# Rapid Parallel Adaptation to Anthropogenic Heavy Metal Pollution

**DOI:** 10.1101/2020.08.12.248328

**Authors:** Alexander S.T. Papadopulos, Andrew J. Helmstetter, Owen G. Osborne, Aaron A. Comeault, Daniel P. Wood, Edward A. Straw, Laurence Mason, Michael F. Fay, Joe Parker, Luke T. Dunning, Andrew D. Foote, Rhian J. Smith, Jackie Lighten

## Abstract

The impact of human mediated environmental change on the evolutionary trajectories of wild organisms is poorly understood. In particular, species’ capacity to adapt rapidly (in hundreds of generations or less), reproducibly and predictably to extreme environmental change is unclear. *Silene uniflora* is predominantly a coastal species, but it has also colonised isolated, disused mines with phytotoxic, zinc-contaminated soils. Here, we found that rapid parallel adaptation to anthropogenic pollution has taken place without geneflow spreading adaptive alleles between populations of the mine ecotype. Across replicate ecotype pairs, we identified shared targets of selection with functions linked to physiological differences between the ecotypes, although the genetic response is only partially shared between mine populations. Our results are consistent with a complex, polygenic genetic architecture underpinning rapid adaptation. This shows that even under a scenario of strong selection and rapid adaptation, evolutionary responses to human activities may be idiosyncratic at the genetic level and, therefore, difficult to predict from genomic data.

## Introduction

Modification of the natural environment by humans has significant implications for biodiversity (Urban 2015; Ceballos *et al*. 2017; Helmstetter *et al*. 2020). Rapid habitat loss or environmental change can drive species to the brink of extinction, but also presents opportunities for adaptation and speciation (Johnson & Munshi-South 2017; Otto 2018; Ravinet *et al*. 2018; Szulkin, M., Munshi-South, J., & Charmantier 2020). There are numerous examples of rapid adaption to human-modified landscapes or activities (McNeilly & Bradshaw 1968; Antonovics & Bradshaw 1970; Wu & Bradshaw 1972; Macnair 1979; Hof *et al*. 2016; Reid *et al*. 2016; Bosse *et al*. 2017), but there is a relatively limited understanding of the extent to which this process is repeatable and predictable (Bay *et al*. 2018; Fitzpatrick *et al*. 2018; Therkildsen *et al*. 2019; Santangelo *et al*. 2020; Van Etten *et al*. 2020). It is also unclear whether such rapid evolution in the wild is typically facilitated by new mutations or draws on standing genetic variation, although there is increasing evidence for the role of standing variation in adaptation generally (Barrett & Schluter 2008; Thompson *et al*. 2019). In part, this uncertainty is due to the difficulty of identifying the key genetic changes that trigger adaptation to a particular selection pressure and, in some cases, identifying the selection pressures themselves.

A powerful approach for determining the genetic architecture of adaptation is to compare taxa that have independently evolved the same adaptive phenotypes on multiple occasions (Rundle *et al*. 2000; Jones *et al*. 2012; Ravinet *et al*. 2016; Nosil *et al*. 2018). Typically taking place tens of thousands or more years ago, such parallel evolution of ecotypes is well known in animals (Soria-Carrasco *et al*. 2014; Ravinet *et al*. 2016; Schweizer *et al*. 2019; Jacobs *et al*. 2020) with fewer clear examples in plants (Roda *et al*. 2013; Trucchi *et al*. 2017; James *et al*. 2020). As a result, there are very few instances in which this kind of parallel adaptation is thought to have taken place rapidly in response to anthropogenic environmental change (Reid *et al*. 2016; Alves *et al*. 2019; Van Etten *et al*. 2020). A significant complication in discriminating between a single or parallel origin of an ecotype (i.e., populations adapted to a specific habitat) is that local gene flow between divergent ecotypes can obscure the true evolutionary relationships of the populations (Ravinet *et al*. 2016; James *et al*. 2020). Promising cases of rapid parallel adaptation do exist, but few have ruled out the possibility of local gene flow creating the false impression of independent origins (James *et al*. 2020). Evidence of contemporary evolution of different ecotypes in sticklebacks has emerged recently to complement the older known events, providing an example where local gene flow is unlikely (Lescak *et al*. 2015; Marques *et al*. 2016). Similarly, the introduction of myxoma virus to rabbits on separate continents since the 1950s was followed by polygenic parallel adaptation towards viral resistance (Alves *et al*. 2019).

Instances where the same toxic chemicals and contaminants have been repeatedly introduced into the environment by humans in isolated locations can generate novel selection regimes that have the potential to promote parallel adaptation. Strong selection, caused by herbicides, pesticides and heavy metals that contaminate soils and water bodies, is capable of producing extremely rapid adaptive responses (Antonovics & Bradshaw 1970; Wu & Bradshaw 1972; Macnair 1979; Hartley *et al*. 2006; Van Etten *et al*. 2020) and trade-offs (Xie & Klerks 2004), and may be particularly prone to triggering parallel responses as a result (MacPherson & Nuismer 2017). Indeed, there is evidence for rapid parallel adaptation from ‘ancient’ standing genetic variation during adaptation to copper mine contamination in two populations of *Mimulus guttatu*s (Wright *et al*. 2015; Lee & Coop 2017). In the Atlantic killifish, *Fundulus heteroclitus*, tolerance to marine pollution has evolved in four populations (Reid *et al*. 2016). The mutations underlying this resistance have evolved on at least two occasions, but migration between three of the four populations may have contributed to the spread of tolerance (Lee & Coop 2017). Convergent herbicide resistance across species is well documented, but there is more limited support for parallel origins within single species and the spread of resistance by gene flow has been harder to rule out (Van Etten *et al*. 2020).

Here, we present evidence for multiple recent and independent origins of heavy metal tolerance in the predominantly coastal plant *Silene uniflora* (sea campion). In Great Britain and Ireland metal mining activities had largely ceased by the early 20th century, but the legacy of spoil heaps and soils contaminated with heavy metals forms a patchwork of highly localised and drastically altered environments across the landscape (Baker *et al*. 2010). Heavy metals, such as zinc, copper, cadmium and lead, are highly toxic to plants, triggering oxidative stress, inhibition of growth and photosynthesis, and death (Küpper & Andresen 2016). As a result, many of these abandoned sites remain barren for hundreds of years after the mining itself has ceased (Baker 1974; Baker *et al*. 2010). Despite its largely linear coastal distribution, *S. uniflora* has managed to colonise a number of isolated inland mine spoils in various regions of the UK and Ireland – although only a small proportion of the >10,800 non-ferrous mines in Great Britain harbour the species (Baker 1974, 1978; Baker & Dalby 1980; Gill 2018). A common feature of the mines that it inhabits is an elevated level of zinc. Experiments in the 1970s demonstrated that mine populations are more zinc tolerant than coastal populations and the plants exclude zinc from their shoots (Baker 1978). Given the generally coastal distribution and the isolated nature of the colonised mines, we hypothesised that the mine ecotypes have independently adapted from the nearest coastal populations. Across four ecotype pairs, we tested for local adaptation using growth experiments to determine whether mine plants are more tolerant to zinc toxicity than their nearest coastal counterparts. We combined a newly sequenced draft genome with reduced representation genotypes for 216 individuals, conducting population genetic analyses to establish the relationships between the populations and whether the mine populations had evolved independently from the physically closest coastal populations. Finally, we used these data to explore the extent to which evolution of the ecotypes is controlled by a parallel/convergent molecular basis.

## Results and discussion

### Anthropogenic adaptation to heavy metal contamination

Populations of *S. uniflora* were sampled from four derelict mines and the nearest coastal population to each across the UK and Ireland (Fig 1A). The contaminated mine sites all have elevated, toxic levels of zinc in the soil (2,410-48,075ppm) relative to typical coastal and inland values (UK mean = 81.3ppm). Lead levels are also high at all mine sites (>10,000ppm; UK mean = 52.6), but only the South Wales (SWA-M) and Irish (IRE-M) mines are heavily contaminated with copper (>10,000 ppm; UK mean = 20.6 ppm; Baker 1978; Allison 2002; Ross *et al*. 2007; Chouinard 2017). We used root elongation experiments with wild collected seed to determine whether mine populations were more tolerant of zinc and copper than the most geographically proximate coastal population. In all cases, mine populations were significantly more zinc tolerant than the local coastal population (Welch’s t-test, two-sided, p <0.005 for all four pairs; Fig 1B). Deep water culture experiments with cuttings from individuals grown in standard conditions also confirmed that plants from mine populations were more zinc tolerant than coastal populations: i.e., root growth continued in mine plants at 600µM ZnSO4, but not in coastal plants (see Methods). However, only the Irish mine population was significantly more copper tolerant than the respective local coastal population (Welch’s t-test, two-sided, p <0.001, Fig 1C). The lack of clear copper tolerance in SWA-M may be due to the relatively high copper concentration used in the experiment, possibly beyond levels that can be tolerated by this population. It is notable that both ecotypes in all Welsh populations were more copper tolerant than the English populations (Fig 1C), suggesting that SWA-M may be able to cope with high copper levels due to constitutive copper tolerance in Welsh *S. uniflora*. These results corroborate earlier findings of zinc and copper tolerance in mine populations of *S. uniflora* (Baker 1978) and demonstrate that all of the sampled mine populations are locally adapted to zinc contamination. Due to the strong selection that heavy metal toxicity exerts, tolerance can evolve in plants within as little as a single generation if there is sufficient genetic variation (Wu & Bradshaw 1972). Although limited mining activity existed at some of these sites as far back as the bronze age, the most intensive working took place between the 18th and 19th centuries (see Methods) and so it is likely that these anthropogenic mine habitats only became available for colonisation once active excavation ceased at mining sites within the last 250 years (Baker 1974). Therefore, populations of the zinc-tolerant ecotype of *S. uniflora* studied here are likely to have evolved since the 18th century (i.e., < 250 generations).

**Figure 1.**
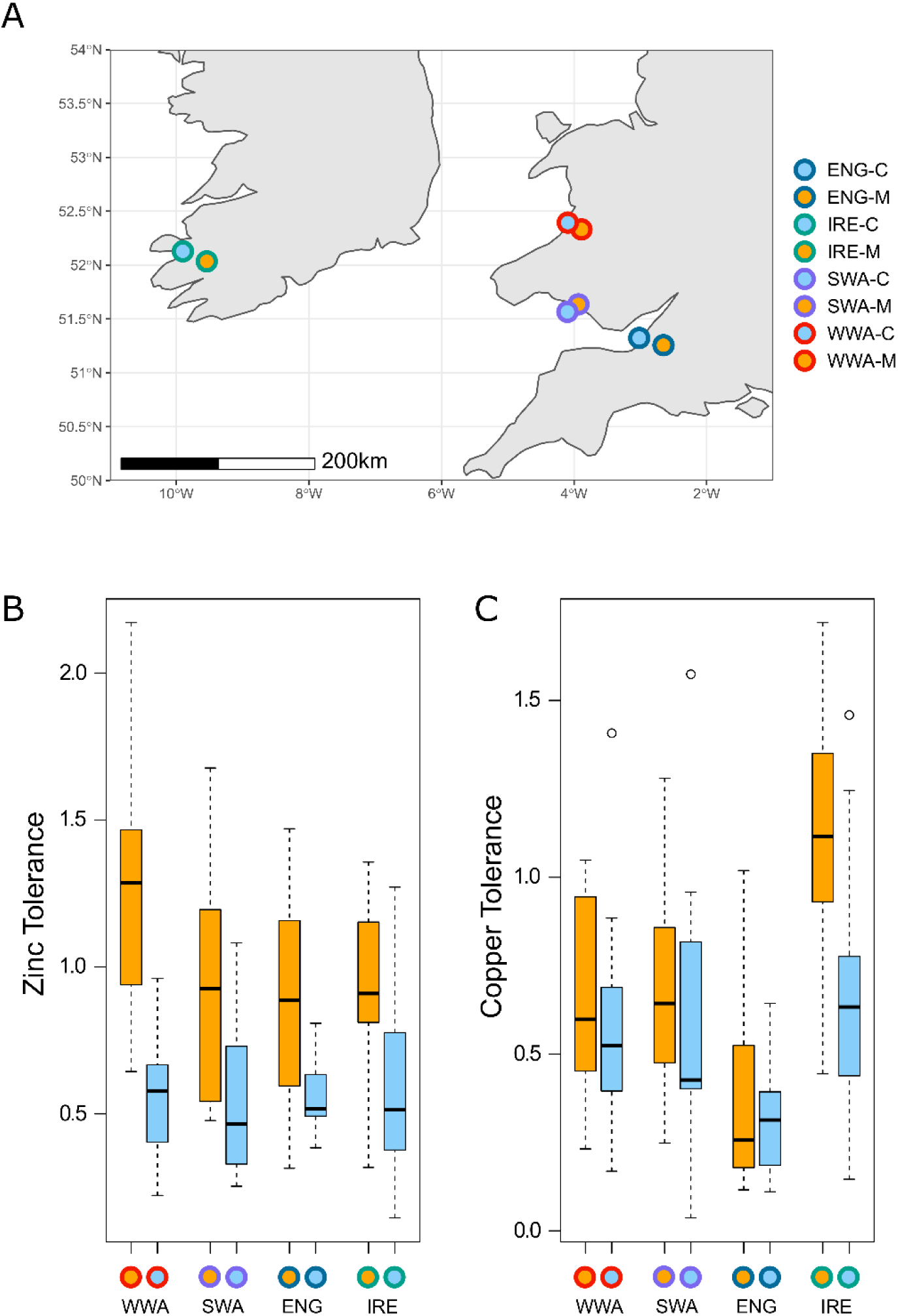
Differential heavy metal tolerance between local mine and coastal ecotypes. **(A)** Map of population sampling locations. Fill colours denote ecotype (mine - orange, coastal - blue). Outline colours denote local populations (West Wales - WWA; South Wales - SWA; South-West England – ENG; South-West Ireland – IRE). The same colour scheme is used throughout. (B) Zinc and (C) copper tolerance for each ecotype pair tolerance (centre line, median; box limits, upper and lower quartiles; whiskers, 1.5x interquartile range; points, outliers; Zinc treatment left to right *n* = 15/17/14/14/14/16/15/19; Copper treatment left to right *n* = 17/17/16/17/15/18/16/18). Local ecotypes have significantly different zinc tolerance, but only the Irish pair have significantly different copper.

### Independent, parallel origins of the mine ecotype

Genetic differentiation between populations was high (mean *F*_ST_ = 0.36; Table S1), reflecting the relatively poor dispersal capabilities and fragmented distribution of the species (Baker 1974; Runyeon & Prentice 1997). Differentiation was substantially higher between mine populations (mean *F*_ST_ = 0.45) than between coastal populations (mean *F*_ST_ = 0.25). Mine ecotypes were also substantially differentiated from the local coastal ecotype (mean *F*_ST_ = 0.36), suggestive of very limited geneflow between ecotypes at the local level. Genetic diversity (π) was also significantly higher in the coastal ecotype versus the mine ecotype (0.065 and 0.044, respectively; Welch’s t-test, two-sided, p <0.036, Table S2). Tajima’s *D* was slightly positive across all populations (mean = 0.24, Table S2), but not significantly different between the ecotypes. As Tajima’s *D* is close to zero, the drop in diversity is unlikely to result from a population bottleneck, but this pattern matches expectations for multiple soft selective sweeps taking place across the genome (Pennings & Hermisson 2006) - as might be expected when colonising a new environment in the face of a strong selection pressure with limited time for new adaptive mutations to evolve.

In the context of recent colonisation, relatively high differentiation and limited gene flow between populations and ecotypes, we predicted that different colonisation scenarios would produce differing patterns of within ecotype isolation by distance (IBD; Wright 1943) - specifically that a scenario of independent origins of the mine ecotype would be distinguishable from a single origin. In a multiple origin scenario, IBD among mine populations should be accentuated relative to the pattern across coastal populations, whereas, in a single origin scenario, IBD within the mine ecotype should be minimal. To test these predictions, we conducted forward-in-time simulations in SLiM v3 (Haller & Messer 2019) and estimated within-ecotype IBD under ‘multiple-origin’ and ‘single-origin’ colonization scenarios (Fig 2A & B, See Methods). As expected, the strength of IBD was significantly higher within the mine ecotype than within the coastal ecotype for the multiple origin scenario (paired t-test, two-sided, p < 0.001; Fig 2A) and the reverse was true for the single origin scenario (paired t-test, two-sided, p < 0.001; Fig 2B). The observed IBD in the sampled populations (Fig 2C) closely matches the expectations for a parallel origin of ecotypes (a similar pattern has been observed in the parallel evolution of Senecio lautus ecotypes; James *et al*. 2020), supporting the hypothesis that the mine habitat has been colonised independently.

**Figure 2.**
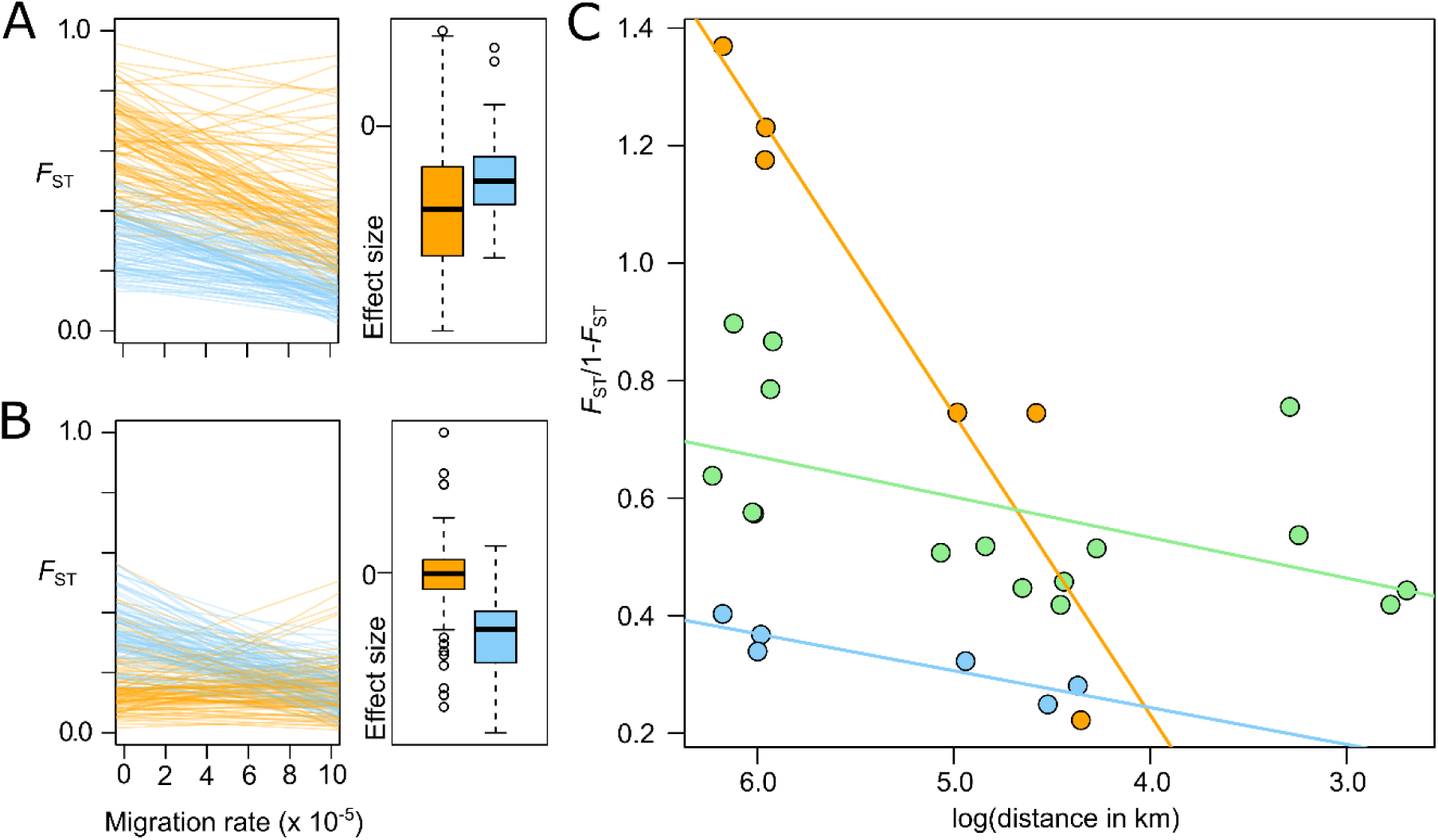
Isolation by distance (IBD) patterns arising from multiple or single origin of the mine ecotype. (A) Under a simulated multiple independent origin model, the correlation between *F*_ST_ and migration between mine populations (orange) is steeper (i.e., IBD is stronger) and has a higher intercept than isolation by distance between coastal populations (blue). (B) In contrast, under a single origin model the relationship between genetic differentiation and geography breaks down between mine populations – the slope is not significantly different from zero and the intercept is lower than between coastal populations (centre line, median; box limits, upper and lower quartiles; whiskers, 1.5x interquartile range; points, outliers). (C) The observed IBD relationships in *S. uniflora* conform to the patterns expected from multiple origins of the mine ecotype. IBD between mine and coastal ecotypes in green.

Phylogenetic reconstruction of evolutionary relationships between the *S. uniflora* populations based on 7,037 linkage disequilibrium pruned SNPs (Fig 3A) and principal component analysis (PCA) of genetic structure from the full set of 74,064 genome aligned SNPs (Fig 3B), clearly indicate three independent origins of the zinc-tolerant mine ecotype; one in Ireland, one in England and one in Wales. The PCA highlights the much higher genetic similarity between coastal populations than between mine populations, which occupy extremely divergent areas of genotype space, suggesting that they may be on different evolutionary trajectories at the genetic level, despite adapting to similar selection pressures. The two Welsh mines are genetically similar (Fig 3B and S1) and, although we cannot rule out independent origins from unsampled non-tolerant populations, it is likely that transport of workers or machinery between Welsh mines dispersed this ecotype between sites. In fact, records of mine ownership from 1758 indicate that human-mediated dispersal is possible between the West Wales and Swansea mines (Hughes 2000). There are at least 14 further records of *S. uniflora* growing on contaminated mine spoil in the UK and Ireland (pers. Obs. & Baker 1974), so our discovery of three independent origins is likely to be a lower bound on the true number of independent origins for the mine ecotype.

**Figure 3.**
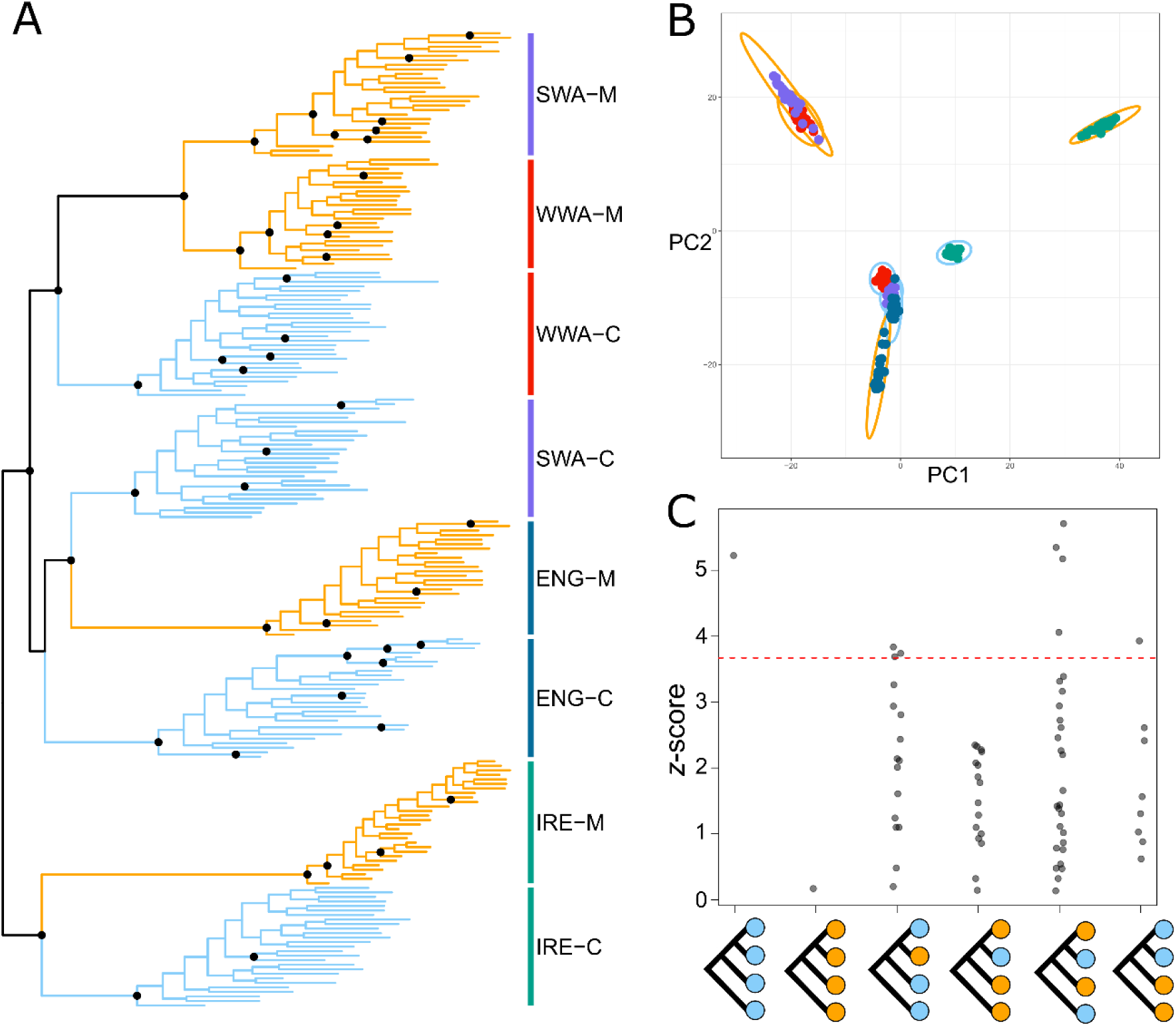
Evidence for three independent origins of the zinc tolerant ecotype in *S. uniflora*. (A) Phylogenetic reconstruction (mine populations in orange and coastal populations in blue). Nodes with greater than 90% bootstrap support are denoted by black circles. (B) Principal components analysis points to three, well supported, independent origins of the zinc tolerant ecotype. Variance explained by PC1 = 12.3% and PC2 = 9.0%. Ecotype and location colours as Fig. 1. (C) *f*4 statistics for the different types of four taxon trees with different combinations of mine and coastal populations. The red line denotes significant deviation from zero after Dunn-Bonferroni correction for multiple tests. There is clear evidence of geneflow from the four coast tree and three coast : one mine trees. On the other hand the four mine and three mine : one coast trees demonstrate that there has not been introgression between mine sites.

Three origins of the mine ecotype were further supported when modelling shared genetic drift among populations (Treemix analysis; Fig S2). This analysis also provided evidence of migration between the Welsh coastal populations (WWA-C and SWA-C) and very weak migration between the Irish, English and Welsh populations. To assess the significance of admixture in the evolution of the mine ecotype, we examined *f*4 statistics across all population quartets (Fig 3C). The *f*4 statistic quantifies shared drift between pairs of populations - significant deviation from an *f*4 statistic of zero for the tree describing the true topology demonstrates that the relationships are not tree-like and is indicative of admixture (Foote & Morin 2016; Lipson 2020). The tree of all four mine populations did not deviate from zero (i.e., there has been no admixture between mines), whereas *f*4 for the coastal population quartet is significantly different from zero - demonstrating that admixture between coastal populations has taken place. Comparisons of quartets with three mine populations and one coastal population provide an additional test of the independent origins of the mine populations, in each case demonstrating that there was no correlated drift between the mine outgroups and the ecotype pair of more closely related populations. On the other hand, the three coastal : one mine comparisons provide further confirmation of geneflow from coastal outgroups into more closely related ecotype pairs in three quartets and support the significance of migration edges between SWA-C and WWA-C, and IRE-C and ENG-C. Overall, our results provide firm support for recent parallel evolution of mine ecotypes, with migration restricted to coastal populations.

### Evidence for molecular convergence/parallelism

To investigate the genetic basis of ecotype differentiation and degree of molecular convergence in adaptation, we conducted pairwise *F*_ST_ -based genome scans for each mine-coast ecotype pair and identified outlier loci potentially under divergent selection. Due to the relatively sparse sampling of our ddRAD dataset and the highly fragmented draft genome (Table S3; N50 = 4,660bp, length = 0.77Gb), we designated genomic scaffolds containing at least one outlier SNP as an outlier scaffold for each comparison (the number of outlier SNPs was not significantly associated with scaffold length; Tukey’s test Fig S3). Across the ecotype pairs, the number of outlier scaffolds ranged from 779-1,216 and the number of outlier SNPs varied from 1,346-2,261 - the degree of overlap between all sets of outlier scaffolds (Fig 4A) and SNPs (Fig 4B) was significantly higher than expected by chance as assessed by Super Exact Test (an extension of Fisher’s Exact Test for multiple sets; Wang *et al*. 2015). In total, 34 scaffolds were identified as outliers across all pairwise ecotype comparisons, while 187 and 756 outlier scaffolds were found across the sets of three and two comparisons, respectively (Fig 4A). There was substantially less overlap at the level of SNPs (Fig 4B), with four shared across all four sets, 85 shared by three sets and 870 shared by two sets. This pattern suggests a highly polygenic basis to mine-coast ecotype differentiation, with a substantial proportion of shared targets of selection found in three or fewer ecotype pairs. However, we are unable to rule out the possibility that the shared scaffolds are physically close to each other in the genome, although linkage disequilibrium between the scaffolds is low (mean r^2^ = 0.021). Dramatically greater overlap in the two Welsh comparisons (WWA and SWA) and a bias towards shared outlier SNPs rather than scaffolds, further supports the single origin of the Welsh mine populations and provides a clear contrast with the degree of outlier overlap with mine populations that evolved in other regions. It is possible that the difference in distribution of overlap between scaffolds and SNPs is due to a limited role of parallelism at the level of individual nucleotides, but greater convergence at the genic level (Conte *et al*. 2012). However, the sparse sampling inherent to the ddRAD approach may mean that the specific adaptive sites are not captured in the analysis (Lowry *et al*. 2017) and there may be more substantial sharing and parallelism of adaptive SNPs across independently evolved mine populations.

**Figure 4.**
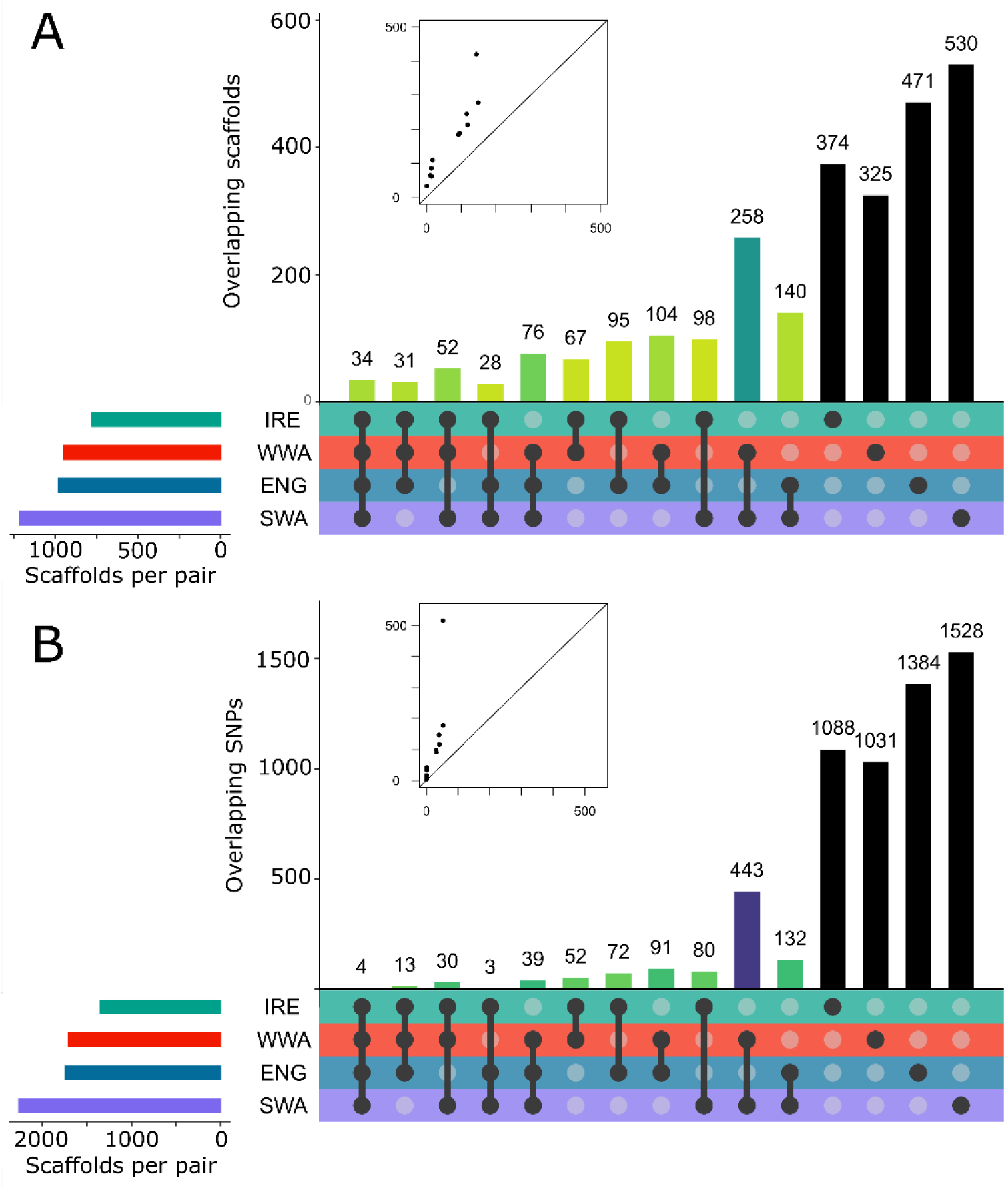
Molecular convergence and divergence across regional ecotype pairs. Upset plots of the shared (A) outlier scaffolds and (B) individual SNPs across the four regional ecotype pairs. Filled points below bars denote which regional sets are intersected for each bar. Inset boxplots show observed overlap (y-axis) vs expected overlap (x-axis) across combinations of regional sets, with line at 1:1. Bars are coloured by super exact test p-value (all < 0.001); darker colours denote smaller p-values.

A polygenic basis to ecotype differentiation in *S. uniflora* is at odds with previous investigations of heavy metal tolerance in *Silene*. Using controlled crosses and hydroponic experiments, these studies indicated that both zinc and copper tolerance have relatively simple genetic bases and are not controlled by the same molecular mechanisms (Schat *et al*. 1996; Schat & Vooijs 1997). The simple architecture for copper tolerance in *S. vulgaris* is also supported by the recent discovery of two related ATPase copper transporters which additively contribute to copper tolerance (Li *et al*. 2017). The potential for polygenic convergence of the mine ecotype in *S. uniflora* is further supported by gene ontology enrichment analysis of the subset of genes found on the 34 scaffolds which were outliers in all 4 pairwise comparisons. This group was significantly enriched for genes involved in metabolism of reactive oxygen species and the regulation of salicylic acid (Table S4), which are critical in responses to cold, salt, drought and heavy metal stresses (Khan *et al*. 2015). Further systematic investigation of gene functions revealed that 15 genes have well supported roles in processes that are relevant to differentiation between coastal and mine plants: eight associated with salt stress, eight with heavy metal stress and four with root development and morphology (Table S5). This points to a potential trade-off in the molecular processes which govern mine-coast ecotype differentiation, with selection against salt-tolerance alleles in mines and against metal tolerance alleles in coastal environments. Alternatively, some alleles for genes that contribute to metal tolerance may be conditionally neutral in coastal plants and under positive selection in the mine environment. In this latter scenario, we might expect a higher incidence of metal tolerance among coastal population, but further work is needed to establish which model underlies local adaptation.

The exact mechanism of zinc tolerance in *Silene* is not well understood. However, hydroponic experiments with mine and coastal ecotypes of *S. uniflora* demonstrated that mine plants grown in zinc-contaminated media accumulate a higher proportion of absorbed zinc in the roots relative to their shoots whereas the reverse is true for coastal plants (Baker 1978). Additional research in *S. vulgaris* indicates that zinc uptake into tonoplast vesicles of zinc-tolerant *S. vulgaris* is higher than in non-tolerant plants (Chardonnens *et al*. 1999). In our study, three genes on outlier scaffolds (*PSD2, WRKY23* and *RWP1*) have direct links to these physiological differences between tolerant and non-tolerant *Silene*: (i) *PSD2* encodes a form of phosphatidylserine decarboxylase which is located in the tonoplast (Nerlich *et al*. 2007), confers cadmium tolerance in *Saccharomyces cerevisiae* (Gulshan *et al*. 2009) and produces phosphatidylethanolamine, which is involved in zinc homeostasis in *Pseudomonas fluorescens* (Appanna *et al*. 1995); (ii) *WRKY23* is a transcription factor that regulates root development by altering auxin distribution through the control of flavanol biosynthesis in *Arabidopsis thaliana* - overexpression of *WRKY23* increases quercetin root concentrations (Grunewald *et al*. 2012). Quercetin is a very efficient chelator of heavy metals (i.e., a molecule that binds metal ions) and supplementation of wild type *A. thaliana* with quercetin stimulates root growth in the presence of zinc ions (Keilig & Ludwig-Müller 2009); and (iii) *RWP1* is required for the production of the cell wall polymer suberin. In *A. thaliana, RWP1* mutants lack suberin and have increased root permeability for NaCl (Gou *et al*. 2009). Furthermore, *Esb1* mutants have increased levels of root suberin, which both decreases accumulation of cadmium, manganese and zinc in the shoots and increases accumulation of sodium in the shoots (Baxter *et al*. 2009). Parallel evolution is expected to be facilitated in spatially structured environments when loci have large, spatially antagonistic fitness effects (Chevin *et al*. 2010). Evidence of such trade-offs in wild plants is lacking, with loci displaying conditional neutrality more commonly detected (Lowry *et al*. 2009; Hall *et al*. 2010; Anderson *et al*. 2011). The dual effect of high suberin levels on restriction of zinc ions to the roots and exposure of the shoots to sodium raises the possibility of a direct trade-off for the ecotypes in suberin production and opens the possibility of antagonistic pleiotropy at *RWP1* influencing the parallel evolution of zinc tolerant ecotypes. Of the three genes, only the scaffold containing RWP1 had consistently lower genetic diversity in the mine ecotype (paired t-test, two-sided, p = 0.030), whereas for *WRKY23* and *PSD2* diversity was only lower in the mines from West Wales and Ireland (Table S2). These findings further support the polygenic nature of parallel adaptation in *S. uniflora* and the potential importance of anatagonistic pleiotropy in the rapid evolution of replicated ecotypes.

In a rapidly changing world, the adaptability of species will be critical for their long-term persistence. This study shows that some species will be capable of responding quickly to strong selection pressures across their range. We argue that plant species with sufficient genetic variation may adapt quickly to a single physiological stress repeatedly in different places, while using subtly different genetic mechanisms. As in *S. uniflora*, those species that evolved to survive in environments with natural sources of high abiotic stress, but which do not compete well in low-abiotic stress/high-biotic competition environments, may be particularly well suited to cope with the ongoing human modification of the planet. Alongside evidence of widespread local adaptation to different environmental conditions in other species (Fournier-Level *et al*. 2011; Papadopulos *et al*. 2014), our findings indicate that while it may be possible to predict which species will adapt to specific environments, the underlying genetic basis to that adaptation may be considerably more variable than is currently understood from the limited number of well-studied examples (Oomen *et al*. 2020). In order to be accurate, predictions of evolutionary responses to environmental change from genomic data will need to account for the possibility that multiple genetic architectures can produce similar phenotypic responses.

## Methods

### Sample collection

Four focal mine sites where *S. uniflora* was known to occur were selected for sampling; Grogwynion (West Wales; WWA; worked 1588 – 1889 C.E.; Hartley 2009), White Rock (Swansea, South Wales; SWA; 1736 – 1928; Hughes 2000), Priddy Pools (Somerset, South-West England; ENG; 1850 – 1908, evidence of Roman mining; Gough 1967) and Ross Island (Co. Kerry, South-West Ireland; IRE; 1707-1829, evidence of Bronze Age mining; O’Brien 2020). For three of these sites (WWA, ENG and IRE), metal tolerance has previously been tested (Baker 1978; Schat *et al*. 1996). White Rock was also located near a previously tested population in Morriston, Swansea (Baker 1978) that no longer exists. The BSBI Database was used to identify the nearest accessible coastal populations to each mine. See Table S6 for population co-ordinates. At each of the eight populations, leaf tissue was sampled from 30 individuals and preserved for DNA extraction in fine mesh silica gel. Individuals were sampled at least one meter apart and samples were collected at even intervals across the extent of each population. At each site, we collected seeds from a minimum of twelve individuals, which were dried and stored separately with silica gel. For assembly of a draft genome, cuttings from a single coastal individual were collected in Tresaith (West Wales), propagated and self-fertilised to produce an inbred F1 (SUTF1P) with reduced heterozygosity.

### Phenotyping

Root elongation experiments were conducted to determine the level of zinc and copper tolerance in each population(Baker 1978). Seeds were germinated in groups of eight (one seed per population) on ¼x Murashige-Skoog media in 1% Agar with no supplemental heavy metals (control treatment), 24µM copper sulphate (copper treatment) or 459 µM zinc sulphate (zinc treatment). Twenty graduated plates were prepared per treatment and the positions of populations within plates was determined using a random seed. Plates were placed upright in a germination cabinet with a 12-hour light/dark cycle for 10 days and then photographed using a digital camera. Radicle length of all seedlings with emerged cotyledons were measured using ImageJ v1.8.0. Zinc and copper tolerance were calculated as the radicle length in the treatment divided by the mean length in the control for each population. Six individuals per population germinated on control media were grown into adults and zinc tolerance was assessed using deep water culture. To do this, cuttings from each individual were rooted in a mist propagator for two weeks before being transferred to a deep-water culture set up with 1/10x Hoagland’s solution. After acclimatisation for one week; the plant roots were stained using a suspension of activated charcoal and rinsed with ddH20, the solution was refreshed and 600 µM Zinc sulphate was added. After a further two days root growth was inspected by eye - the presence of unstained root tips (i.e., ongoing root growth) was taken as confirmation of zinc tolerance (Schat *et al*. 1996; Bratteler *et al*. 2006b).

### Genome assembly

DNA was extracted from silica dried leaf tissue using Qiagen DNeasy Plant tissue kits. DNA quality was assessed using agarose gel electrophoresis and DNA was quantified using a Promega Quantus fluorometer with Quantifluor dsDNA kits. For draft genome assembly, four NEBnext Ultra II libraries were prepared for SUTF1P and each was sequenced using illumina MiSeq v3 600bp PE cartridges. Adapter and quality trimming were performed using cutadapt v2.1 (Martin 2011) and Trimmomatic v0.36(Bolger *et al*. 2014) (minimum quality = 15, minimum length = 64). Overlapping read pairs were merged using Abyss-mergepairs (Jackman *et al*. 2017) and non-overlapping pairs merged using konnector v2.0 (Vandervalk *et al*. 2015) with a bloom filter containing merged and unmerged reads for all libraries (kmer length=96, bloom filter FPR = 1.01%). illumina reads were assembled into contigs using Abyss v2.0 (Jackman *et al*. 2017) with a kmer length=241 – selected after estimation with kmergenie v1.7048 and Abyss runs with kmers = 96/127/151. To scaffold the assembly the same individual was sequenced using an Oxford Nanopore MinION (Three R9 flow cells and one R9.4 flow cell with SQK-NSK007 kits). Nanopore reads were corrected with Proovread v2.12 (Hackl *et al*. 2014) using the processed illumina reads. Redundans v0.14a (Pryszcz & Gabaldón 2016) was used to reduce contig redundancy caused by heterozygosity (minimum identity 95%) and scaffold contigs using the corrected nanopore data. Abyss-sealer (Paulino *et al*. 2015) was used to fill gaps in the scaffolded assembly (kmers = 94/89/84) and completeness was assessed with BUSCO v3 (Benchmarking Universal Single-Copy Orthologs; complete and fragmented = 78.5%, Table S3). Augustus (Stanke *et al*. 2006) was used to predict genes in the genomic scaffolds using the annotation training files from *Solanum lycopersicum*. The resulting predicted amino acid sequences were BLASTp-searched (Camacho *et al*. 2009) against the *Arabidopsis thaliana* proteome (Araport11) and only the best scoring hit from each predicted amino acid sequence was retained.

### Genotyping

Double-digest RAD sequencing was performed following a modified protocol of Peterson et al (2012) detailed in Papadopulos et al (2019a, b). For this study, size selection was conducted with a pippin prep (468-546bp) and one pool of 230 uniquely barcoded individuals was sequenced on five lanes of an illumina HiSeq 2500 (100bp, PE) at the Earlham Institute. Raw reads were demultiplexed, trimmed to 90bp and low-quality reads were discarded, resulting in an average of 4.76M reads per sample (s.d. 2.01M). Reads were mapped to the draft genome using bowtie v2.3.4 (Langmead & Salzberg 2012) in end-to-end mode and excluding reads with low mapping quality (Q<20). SNPS were called from the resulting BAM files using gstacks v2.0b (Rochette *et al*. 2019), 14 samples were excluded from further analysis due to low coverage. Genotypes for SNPS with less than 20% missing data were extracted in VCF and RADpainter format using Populations v2.0b (Rochette *et al*. 2019). In total, 216 individuals were genotyped at 74,064 SNPs.

### Evolutionary genetics

Population genetic structure across *S. uniflora* was assessed using principal components analysis implemented in adegenet v2.1.3 (Jombart 2008) in R and genetic co-ancestry was estimated using the haplotype-based inference method of fineRADstructure v0.3.2 (Malinsky *et al*. 2018). To assess patterns of isolation by distance, pairwise genetic differentiation between the sampled populations (Weir and Cockerham’s *F*_ST_) was calculated using Arlequin v3.5.2.2 (Excoffier & Lischer 2010), pairwise geographic distances between populations were calculated with the distm function in the geosphere R package and isolation by distance estimated in R using linear regression. Tajima’s *D* was calculated for 20kb sliding windows in VCFtools v0.1.16 (Danecek *et al*. 2011) and averaged over the subset of windows for which *D* could be calculated in all populations. To identify the isolation by distance signature expected from parallel vs single origins of the mine ecotypes, we conducted simulations in SLiM v3.3.2 under two scenarios: Independent colonisation of mines from the nearest coastal population and non-independent colonisation of mines from the same individual coastal population. In the latter case, the ‘founding’ coastal population was randomly chosen in each independent iteration of the simulation. All simulations were initiated with a burn-in period of 100,000 generations and a population size of 10,000 individuals. Each individual in the population was diploid and hermaphroditic, and generations were non-overlapping (i.e. Wright-Fisher simulations). To track genetic relationships among populations, we simulated a single chromosome that was 50,000 bp long with a uniform mutation rate of 7.5 × 10^−9^ - based on estimates for *S. latifolia* (Krasovec *et al*. 2018) - and a recombination rate of 4.0 × 10^−9^ - based on the genetic map length (446cM; Bratteler *et al*. 2006a) and genome size (1.13Gb) of *S. vulgaris* (Pellicer & Leitch 2020). In the 100,000^th^ generation, two populations (p1 and p2) were colonised with 500 individuals each from the ancestral population. These two populations represented those that initially colonised Ireland and the west coast of England/Wales at the end of the Last Glacial Maximum. Subsequent stepwise colonisation of populations (i.e. p2 -> p3 -> p4), representing coastal populations, occurred every 20 generations until there were four coastal populations in the 100,040th generation. Coastal populations were always founded with 500 individuals and population sizes increased to 1,000 individuals ten generations after a population was initially founded. After colonisation, p1 and p2 exchanged migrants at a rate of 0.00001 per generation, p2 and p3 at a rate of 0.0001, and p3 and p4 at a rate of 0.0001. P1 through p4 were therefore effectively arranged along a line and migration rates between non-adjacent populations were equivalent to the product of migration rates connecting them. Ten-thousand generations after the ‘coastal’ populations were founded, 100 individuals were used to found each of four populations meant to reflect those found in mine environments. After founding the ‘mine’ populations, these populations exchanged migrants with the nearest ‘coastal’ population at a rate of 0.0002. All populations then evolved for an additional 100 generations. At the end of the simulations (i.e. at generation 110,150), we calculated and output *F*_ST_ between each of the four ‘coastal’ populations (all pairwise comparisons) and each of the four ‘mine’ populations. We ran 100 independent replicates for each of the three colonisation scenarios described above.

To further establish the evolutionary relationships between the populations, the dataset was pruned to 7,037 SNPs using a linkage disequilibrium threshold of 0.1 and minor allele frequency threshold of 0.05, and the phylogenetic tree estimated with 1,000 bootstrap replicates using the maximum likelihood approach implemented in SNPhylo v2 (Lee *et al*. 2014). This reduced dataset was then used to explore the possibility of migration and introgression between the populations using Treemix v0.1.15 (Pickrell & Pritchard 2012). The maximum likelihood estimation of the tree in Treemix, one to ten migration edges were fitted and the number of edges that explained 99.8% of the variance selected as the best model. Using the fourpop function in Treemix, *f*4 statistics (Reich *et al*. 2009) were calculated for all population quartets to assess whether relationships between the populations deviated significantly (after Dunn-Bonferroni correction) from tree-likeness.

To investigate the level of parallel evolution at the molecular level we calculated Weir and Cockerham’s *F*_ST_ at all variable sites in pairwise comparisons between the geographically proximate ecotype pairs using VCFtools v0.1.16. SNPs falling in the upper 95% percentile of values in each pairwise comparison were designated as outlier loci and scaffolds containing one of more outlier SNPS were designated as outlier scaffolds. Overlap of outlier SNPs and scaffolds was visualised using upsetR v1.4.0 (Conway *et al*. 2017) and significance of overlap was assessed using SuperExactTest v1.0.7 (Wang *et al*. 2015). To investigate the possible functions of genes in outlier regions, all genes on the outlier scaffolds that were in common across the four pairwise ecotype comparisons were subjected to gene ontology enrichment analysis performed in topGo v3.11 (Alexa & Rahnenfuhrer 2020) using the “elim” algorithm and Fisher’s Exact tests to assess significance. Further assessment of gene functions were made from The *Arabidopsis* Information Resource (TAIR) descriptions and associated references. Systematic searches were performed using gene names with and without the terms “stress” and “heavy metal” using Google Scholar.

## Acknowledgements

We thank Natural Environment Research Council (NERC) and the Royal Society for funding; Alan Baker, Roger Butlin, Andrew Leitch & Steve Rossiter for encouragement and discussion; Robyn Cowan, Wendy Grail & Jonathan Kendon for laboratory support; and the Botanical Society of the British Isles & Mike Gill for access to databases.

## Author Contributions

ASTP conceived and designed the research with contributions from all co-authors. ASTP, RJS, JL & ES conducted fieldwork. AJH conducted ddRAD lab work. ASTP, ES & LM conducted tolerance experiments. ASTP analysed the data, with contributions from OGO, AF, AC & JL. AAC conducted simulations. ASTP wrote the manuscript and all authors commented on the final version.

## References

Alexa, A. & Rahnenfuhrer, J. (2020). topGO: Enrichment analysis for gene ontology. R package version 2.40.0.

Allison, S. (2002). Distribution and biogeochemical cycling of trace metals in soils at an abandoned lead mining and smelting. University of the West of England, Bristol.

Alves, J.M., Carneiro, M., Cheng, J.Y., de Matos, A.L., Rahman, M.M., Loog, L., et al. (2019). Parallel adaptation of rabbit populations to myxoma virus. Science., 363, 1319–1326.

Anderson, J.T., Willis, J.H. & Mitchell-Olds, T. (2011). Evolutionary genetics of plant adaptation. Trends Genet., 27, 258–266.

Antonovics, J. & Bradshaw, A.D. (1970). Evolution in closely adjacent plant poplations VIII. Clinal patterns at a mine boundary. Heredity., 25, 349–362.

Appanna, V.D., Finn, H. & Pierre, M.S. (1995). Exocellular phosphatidylethanolamine production and multiple-metal tolerance in *Pseudomonas fluorescens*. FEMS Microbiol. Lett., 131, 53–56.

Baker, A.J.M. (1978). Ecophysiological aspects of zinc tolerance in *Silene maritima* With. New Phytol., 80, 635–642.

Baker, A.J.M. & Dalby, D.H. (1980). Morphological variation between some isolated populations of *Silene maritima* With. in the British Isles with particular reference to inland populations on metalliferous soils. New Phytol., 84, 123–138.

Baker, A.J.M., Ernst, W.H.O., van der Ent, A., Malaisse, F. & Ginocchio, R. (2010). Metallophytes: the unique biological resource, its ecology and conservational status in Europe, central Africa and Latin America. In: Ecology of industrial pollution (eds. Lesley C. Batty & Hallberg, K.B.). pp. 7–39.

Baker, A.J.M.M. (1974). Heavy metal tolerance and population differentiation in *Silene maritima* With. Imperial College London.

Barrett, R.D.H. & Schluter, D. (2008). Adaptation from standing genetic variation. Trends Ecol. Evol., 23, 38–44.

Baxter, I., Hosmani, P.S., Rus, A., Lahner, B., Borevitz, J.O., Muthukumar, B., et al. (2009). Root suberin forms an extracellular barrier that affects water relations and mineral nutrition in *Arabidopsis*. PLoS Genet., 5.

Bay, R., Harrigan, R., Le Underwood, V., Gibbs, H., Smith, T. & K, R. (2018). Genomic signals of selection predict climate-driven population declines in a migratory bird. Science., 361, 83–86.

Bolger, A.M., Lohse, M. & Usadel, B. (2014). Trimmomatic: A flexible trimmer for Illumina sequence data. Bioinformatics, 30, 2114–2120.

Bosse, M., Spurgin, L.G., Laine, V.N., Cole, E.F., Firth, J.A., Gienapp, P., et al. (2017). Recent natural selection causes adaptive evolution of an avian polygenic trait. Science., 358, 365–368.

Bratteler, M., Lexer, C. & Widmer, A. (2006a). A genetic linkage map of *Silene vulgaris* based on AFLP markers. Genome, 49, 320–327.

Bratteler, M., Lexer, C. & Widmer, A. (2006b). Genetic architecture of traits associated with serpentine adaptation of *Silene vulgaris*. J. Evol. Biol., 19, 1149–1156.

Camacho, C., Coulouris, G., Avagyan, V., Ma, N., Papadopoulos, J., Bealer, K., et al. (2009). BLAST plus: architecture and applications. BMC Bioinformatics, 10, 1.

Ceballos, G., Ehrlich, P.R. & Dirzo, R. (2017). Biological annihilation via the ongoing sixth mass extinction signaled by vertebrate population losses and declines. Proc. Natl. Acad. Sci. U.S.A., 114, 6089–6096.

Chardonnens, A.N., Koevoets, P.L.M., van Zanten, A., Schat, H. & Verkleij, J. a. C. (1999). Properties of enhanced tonoplast zinc transport in naturally selected zinc-tolerant *Silene vulgaris*. Plant Physiol., 120, 779–786.

Chevin, L.M., Martin, G. & Lenormand, T. (2010). Fisher’s model and the genomics of adaptation: Restricted pleiotropy, heterogenous mutation, and parallel evolution. Evolution., 64, 3213–3231.

Chouinard, C. (2017). Elemental contamination of an ancient copper mine in Killarney, Ireland. Augustana College, Rock Island Illinois.

Conte, G.L., Arnegard, M.E., Peichel, C.L. & Schluter, D. (2012). The probability of genetic parallelism and convergence in natural populations. Proc. R. Soc. B Biol. Sci., 279, 5039–5047.

Conway, J.R., Lex, A. & Gehlenborg, N. (2017). UpSetR: An R package for the visualization of intersecting sets and their properties. Bioinformatics, 33, 2938–2940.

Danecek, P., Auton, A., Abecasis, G., Albers, C.A., Banks, E., DePristo, M.A., et al. (2011). The variant call format and VCFtools. Bioinformatics, 27, 2156–2158.

Van Etten, M., Lee, K.M., Chang, S.M. & Baucom, R.S. (2020). Parallel and nonparallel genomic responses contribute to herbicide resistance in *Ipomoea purpurea*, a common agricultural weed. PLoS Genet., 16, e1008593.

Excoffier, L. & Lischer, H.E.L. (2010). Arlequin suite ver 3.5: a new series of programs to perform population genetics analyses under Linux and Windows. Mol. Ecol. Resour., 10, 564–567.

Fitzpatrick, M.C., Keller, S.R. & Lotterhos, K.E. (2018). Comment on “Genomic signals of selection predict climate-driven population declines in a migratory bird.” Science., 361, 2–4.

Foote, A.D. & Morin, P.A. (2016). Genome-wide SNP data suggest complex ancestry of sympatric North Pacific killer whale ecotypes. Heredity., 117, 316–325.

Fournier-Level, A., Korte, A., Cooper, M.D., Nordborg, M., Schmitt, J. & Wilczek, A.M. (2011). A map of local adaptation in *Arabidopsis thaliana*. Science., 334, 86–89.

Gill, M. (2018). Gazetteer of British Miscellaneous Mines.

Gou, J.Y., Yu, X.H. & Liu, C.J. (2009). A hydroxycinnamoyltransferase responsible for synthesizing suberin aromatics in *Arabidopsis*. Proc. Natl. Acad. Sci. U. S. A., 106, 18855–18860.

Gough, J.W. (1967). The Mines of Mendip. David & Charles.

Grunewald, W., De Smet, I., Lewis, D.R., Löfke, C., Jansen, L., Goeminne, G., et al. (2012). Transcription factor WRKY23 assists auxin distribution patterns during *Arabidopsis* root development through local control on flavonol biosynthesis. Proc. Natl. Acad. Sci. U. S. A., 109, 1554–1559.

Gulshan, K., Shahi, P. & Moye-Rowley, W.S. (2009). Compartment-specific synthesis of phosphatidylethanolamine is required for normal heavy metal resistance. Mol. Biol. Cell, 21, 443–455.

Hackl, T., Hedrich, R., Schultz, J. & Förster, F. (2014). Proovread: Large-scale high-accuracy PacBio correction through iterative short read consensus. Bioinformatics, 30, 3004–3011.

Hall, M.C., Lowry, D.B. & Willis, J.H. (2010). Is local adaptation in *Mimulus guttatus* caused by trade-offs at individual loci? Mol. Ecol., 19, 2739–2753.

Haller, B.C. & Messer, P.W. (2019). SLiM 3: Forward genetic simulations beyond the Wright-Fisher model. Mol. Biol. Evol., 36, 632–637.

Hartley, C.J., Newcomb, R.D., Russell, R.J., Yong, C.G., Stevens, J.R., Yeates, D.K., et al. (2006). Amplification of DNA from preserved specimens shows blowflies were preadapted for the rapid evolution of insecticide resistance. Proc. Natl. Acad. Sci. U. S. A., 103, 8757–8762.

Hartley, S. (2009). Remediation of abandoned metal mine drainage using dealginated seaweed.

Helmstetter, A.J., Cable, S., Rakotonasolo, F., Rabarijaona, R., Rakotoarinivo, M., Eiserhardt, W.L., et al. (2020). The demographic history of micro-endemics: have rare species always been rare? bioRxiv.

Hof, A.E.V. t., Campagne, P., Rigden, D.J., Yung, C.J., Lingley, J., Quail, M.A., et al. (2016). The industrial melanism mutation in British peppered moths is a transposable element. Nature, 534, 102–105.

Hughes, S. (2000). Copperopolis: Landscapes of the Early Industrial Period in Swansea. Royal Commission on the Ancient and Historical Monuments of Wales.

Jackman, S.D., Vandervalk, B.P., Mohamadi, H., Chu, J., Yeo, S., Hammond, S.A., et al. (2017). ABySS 2.0: Resource-efficient assembly of large genomes using a Bloom filter. Genome Res., 27, 768–777.

Jacobs, A., Carruthers, M., Yurchenko, A., Gordeeva, N. V., Alekseyev, S.S., Hooker, O., et al. (2020). Parallelism in eco-morphology and gene expression despite variable evolutionary and genomic backgrounds in a Holarctic fish. PLOS Genet., 16, e1008658.

James, M.E., Arenas-Castro, H., Groh, J.S., Engelstaedter, J. & Ortiz-Barrientos, D. (2020). Highly replicated evolution of parapatric ecotypes. bioRxiv, 2020.02.05.936401.

Johnson, M.T.J. & Munshi-South, J. (2017). Evolution of life in urban environments. Science., 358.

Jombart, T. (2008). Adegenet: A R package for the multivariate analysis of genetic markers. Bioinformatics, 24, 1403–1405.

Jones, F.C., Grabherr, M.G., Chan, Y.F., Russell, P., Mauceli, E., Johnson, J., et al. (2012). The genomic basis of adaptive evolution in threespine sticklebacks. Nature, 484, 55–61.

Keilig, K. & Ludwig-Müller, J. (2009). Effect of flavonoids on heavy metal tolerance in *Arabidopsis thaliana* seedlings. Bot. Stud., 50, 311–318.

Khan, M.I.R., Fatma, M., Per, T.S., Anjum, N.A. & Khan, N.A. (2015). Salicylic acid-induced abiotic stress tolerance and underlying mechanisms in plants. Front. Plant Sci., 6, 1–17.

Krasovec, M., Chester, M., Ridout, K. & Filatov, D.A. (2018). The mutation rate and the age of the sex chromosomes in *Silene latifolia*. Curr. Biol., 28, 1832-1838.e4.

Küpper, H. & Andresen, E. (2016). Mechanisms of metal toxicity in plants. Metallomics, 8, 269–285.

Langmead, B. & Salzberg, S.L. (2012). Fast gapped-read alignment with Bowtie 2. Nat. Methods, 9, 357–359.

Lee, K.M. & Coop, G. (2017). Distinguishing among modes of convergent adaptation using population genomic data. Genetics, 207, 1591–1619.

Lee, T.H., Guo, H., Wang, X., Kim, C. & Paterson, A.H. (2014). SNPhylo: A pipeline to construct a phylogenetic tree from huge SNP data. BMC Genomics, 15, 1–6.

Lescak, E.A., Bassham, S.L., Catchen, J., Gelmond, O., Sherbick, M.L., von Hippel, F.A., et al. (2015). Evolution of stickleback in 50 years on earthquake-uplifted islands. Proc. Natl. Acad. Sci. U.S.A., 201512020.

Li, Y., Iqbal, M., Zhang, Q., Spelt, C., Bliek, M., Hakvoort, H.W.J., et al. (2017). Two *Silene vulgaris* copper transporters residing in different cellular compartments confer copper hypertolerance by distinct mechanisms when expressed in *Arabidopsis thaliana*. New Phytol., 215, 1102–1114.

Lipson, M. (2020). Interpreting f-statistics and admixture graphs : theory and examples. Preprints, 1–18.

Lowry, D.B., Hall, M.C., Salt, D.E. & Willis, J.H. (2009). Genetic and physiological basis of adaptive salt tolerance divergence between coastal and inland *Mimulus guttatus*. New Phytol., 183, 776–788.

Lowry, D.B., Hoban, S., Kelley, J.L., Lotterhos, K.E., Reed, L.K., Antolin, M.F., et al. (2017). Breaking RAD: an evaluation of the utility of restriction site-associated DNA sequencing for genome scans of adaptation. Mol. Ecol. Resour., 17, 142–152.

Macnair, M.R. (1979). The genetics of copper tolerance in the yellow monkey flower, *Mimulus gutattus*. I. Crosses to nontolerants. Genetics, 91, 553–563.

MacPherson, A. & Nuismer, S.L. (2017). The probability of parallel genetic evolution from standing genetic variation. J. Evol. Biol., 30, 326–337.

Malinsky, M., Trucchi, E., Lawson, D.J. & Falush, D. (2018). RADpainter and fineRADstructure: Population Inference from RADseq Data. Mol. Biol. Evol., 35, 1284–1290.

Marques, D.A., Lucek, K., Meier, J.I., Mwaiko, S., Wagner, C.E., Excoffier, L., et al. (2016). Genomics of rapid incipient speciation in sympatric threespine stickleback. PLoS Genet., 12, 1–34.

Martin, M. (2011). Cutadapt removes adapter sequences from high-throughput sequencing reads, EMBnet.journal 17. EMBnet.journal, 17.

McNeilly, T. & Bradshaw, A.D. (1968). Evolutionary processes in populations of copper tolerant *Agrostis tenuis*. Evolution (N. Y)., 22, 108–118.

Nerlich, A., Von Orlow, M., Rontein, D., Hanson, A.D. & Dörmann, P. (2007). Deficiency in phosphatidylserine decarboxylase activity in the psd1 psd2 psd3 triple mutant of *Arabidopsis* affects phosphatidylethanolamine accumulation in mitochondria. Plant Physiol., 144, 904–914.

Nosil, P., Villoutreix, R., De Carvalho, C.F., Farkas, T.E., Soria-Carrasco, V., Feder, J.L., et al. (2018). Natural selection and the predictability of evolution in *Timema* stick insects. Science., 359, 765–770.

O’Brien, W. (2020). Ross Island Mine Heritage. Dep. Archaeol. NUI Galw. Available at: http://www.nuigalway.ie/ross_island/index.htm. Last accessed 13 May 2020.

Oomen, R.A., Kuparinen, A. & Hutchings, J.A. (2020). Consequences of single-locus and tightly linked genomic architectures for evolutionary responses to environmental change. J. Hered., esaa020.

Otto, S.P. (2018). Adaptation, speciation and extinction in the Anthropocene. Proc. R. Soc. B Biol. Sci., 285.

Papadopulos, A.S.T., Igea, J., Dunning, L.T., Osborne, O.G., Quan, X., Pellicer, J., et al. (2019a). Ecological speciation in sympatric palms: 3. Genetic map reveals genomic islands underlying species divergence in *Howea*. Evolution (N. Y)., 73, 1996–2002.

Papadopulos, A.S.T., Igea, J., Smith, T.P., Hutton, I., Baker, W.J., Butlin, R.K., et al. (2019b). Ecological speciation in sympatric palms: 4. Demographic analyses support speciation of *Howea* in the face of high gene flow. Evolution (N. Y)., 73, 1996–2002.

Papadopulos, A.S.T., Kaye, M., Devaux, C., Hipperson, H., Lighten, J., Dunning, L.T., et al. (2014). Evaluation of genetic isolation within an island flora reveals unusually widespread local adaptation and supports sympatric speciation. Philos. Trans. R. Soc. B Biol. Sci., 369, 20130342.

Paulino, D., Warren, R.L., Vandervalk, B.P., Raymond, A., Jackman, S.D. & Birol, I. (2015). Sealer: A scalable gap-closing application for finishing draft genomes. BMC Bioinformatics, 16, 1–8.

Pellicer, J. & Leitch, I.J. (2020). The Plant DNA C-values database (release 7.1): an updated online repository of plant genome size data for comparative studies. New Phytol., 226, 301–305.

Pennings, P.S. & Hermisson, J. (2006). Soft sweeps III: The signature of positive selection from recurrent mutation. PLoS Genet., 2, 1998–2012.

Peterson, B.K., Weber, J.N., Kay, E.H., Fisher, H.S. & Hoekstra, H.E. (2012). Double digest RADseq: an inexpensive method for de novo SNP discovery and genotyping in model and non-model species. PLoS One, 7, e37135.

Pickrell, J.K. & Pritchard, J.K. (2012). Inference of population splits and mixtures from genome-wide allele frequency data. PLoS Genet., 8.

Pryszcz, L.P. & Gabaldón, T. (2016). Redundans: An assembly pipeline for highly heterozygous genomes. Nucleic Acids Res., 44, e113.

Ravinet, M., Elgvin, T.O., Trier, C., Aliabadian, M., Gavrilov, A. & Sætre, G.P. (2018). Signatures of human-commensalism in the house sparrow genome. Proc. R. Soc. B Biol. Sci., 285.

Ravinet, M., Westram, A., Johannesson, K., Butlin, R., André, C. & Panova, M. (2016). Shared and nonshared genomic divergence in parallel ecotypes of Littorina saxatilis at a local scale. Mol. Ecol., 25, 287–305.

Reich, D., Thangaraj, K., Patterson, N., Price, A.L. & Singh, L. (2009). Reconstructing Indian population history. Nature, 461, 489–494.

Reid, N.M., Proestou, D.A., Clark, B.W., Warren, W.C., Colbourne, J.K., Shaw, J.R., et al. (2016). The genomic landscape of rapid repeated evolutionary adaptation to toxic pollution in wild fish. Science., 354, 1305–1308.

Rochette, N.C., Rivera-Colón, A.G. & Catchen, J.M. (2019). Stacks 2: Analytical methods for paired-end sequencing improve RADseq-based population genomics. Mol. Ecol., 28, 4737–4754.

Roda, F., Ambrose, L., Walter, G.M., Liu, H.L., Schaul, A., Lowe, A., et al. (2013). Genomic evidence for the parallel evolution of coastal forms in the *Senecio lautus* complex. Mol. Ecol., 22, 2941–2952.

Ross, S.M., Wood, M.D., Copplestone, D., Warriner, M. & Crook, P. (2007). UK Soil and Herbage - Pollutant Survey Environmental Concentrations of Heavy Metals in UK Soil and Herbage.

Rundle, H.D., Nagel, L., Wenrick Boughman, J. & Schluter, D. (2000). Natural selection and parallel speciation in sympatric sticklebacks. Science., 287, 306–308.

Runyeon, H. & Prentice, H.C. (1997). Genetic differentiation in the Bladder campions, *Silene vulgaris* and *S. uniflora* (Caryophyllaceae), in Sweden. Biol. J. Linn. Soc., 61, 559–584.

Santangelo, J.S., Thompson, K.A., Cohan, B., Syed, J., Ness, R.W. & Johnson, M.T.J. (2020). Predicting the strength of urban-rural clines in a Mendelian polymorphism along a latitudinal gradient. Evol. Lett., 4, 212–225.

Schat, H. & Vooijs, R. (1997). Multiple tolerance and co-tolerance to heavy metals in *Silene vulgaris*: A co-segregation analysis. New Phytol., 136, 489–496.

Schat, H., Vooijs, R. & Kuiper, E. (1996). Identical major gene loci for heavy metal tolerances that have independently evolved in different local populations and subspecies of *Silene vulgaris*. Evolution., 50, 1888–1895.

Schweizer, M., Warmuth, V., Alaei Kakhki, N., Aliabadian, M., Förschler, M., Shirihai, H., et al. (2019). Parallel plumage colour evolution and introgressive hybridization in wheatears. J. Evol. Biol., 32, 100–110.

Soria-Carrasco, V., Gompert, Z., Comeault, A.A., Farkas, T.E., Parchman, T.L., Johnston, J.S., et al. (2014). Stick insect genomes reveal natural selection’s role in parallel speciation. Science., 344, 738–742.

Stanke, M., Schöffmann, O., Morgenstern, B. & Waack, S. (2006). Gene prediction in eukaryotes with a generalized hidden Markov model that uses hints from external sources. BMC Bioinformatics, 7, 62.

Szulkin, M., Munshi-South, J., & Charmantier, A. (2020). Urban Evolutionary Biology. Oxford University Press.

Therkildsen, N.O., Wilder, A.P., Conover, D.O., Munch, S.B., Baumann, H. & Palumbi, S.R. (2019). Contrasting genomic shifts underlie parallel phenotypic evolution in response to fishing. Science., 490, 487–490.

Thompson, K.A., Osmond, M.M. & Schluter, D. (2019). Parallel genetic evolution and speciation from standing variation. Evol. Lett., 3, 129–141.

Trucchi, E., Frajman, B., Haverkamp, T.H.A., Schönswetter, P. & Paun, O. (2017). Genomic analyses suggest parallel ecological divergence in *Heliosperma pusillum* (Caryophyllaceae). New Phytol., 216, 267–278.

Urban, M. (2015). Accelerating extinction risk from climate change. Science., 348.

Vandervalk, B.P., Yang, C., Xue, Z., Raghavan, K., Chu, J., Mohamadi, H., et al. (2015). Konnector v2.0: Pseudo-long reads from paired-end sequencing data. BMC Med. Genomics, 8, S1.

Wang, M., Zhao, Y. & Zhang, B. (2015). Efficient test and visualization of multi-set intersections. Sci. Rep., 5, 1–12.

Wright, K.M., Hellsten, U., Xu, C., Jeong, A.L., Sreedasyam, A., Chapman, J.A., et al. (2015). Adaptation to heavy-metal contaminated environments proceeds via selection on pre-existing genetic variation. bioRxiv, 029900.

Wright, S. (1943). Isolation by distance. Genetics, 28, 114.

Wu, L. & Bradshaw, A.D. (1972). Aerial pollution and the rapid evolution of copper tolerance. Nature, 238, 167–169.

Xie, L. & Klerks, P.L. (2004). Fitness cost of resistance to cadmium in the least killifish (*Heterandria formosa*). Environ. Toxicol. Chem., 23, 1499–1503.

