## Supplementary Information for "Rapid Parallel Adaptation to Anthropogenic Heavy Metal Pollution"

**Figure S1.** Co-ancestry matrix from finRADstructure clustered by population.

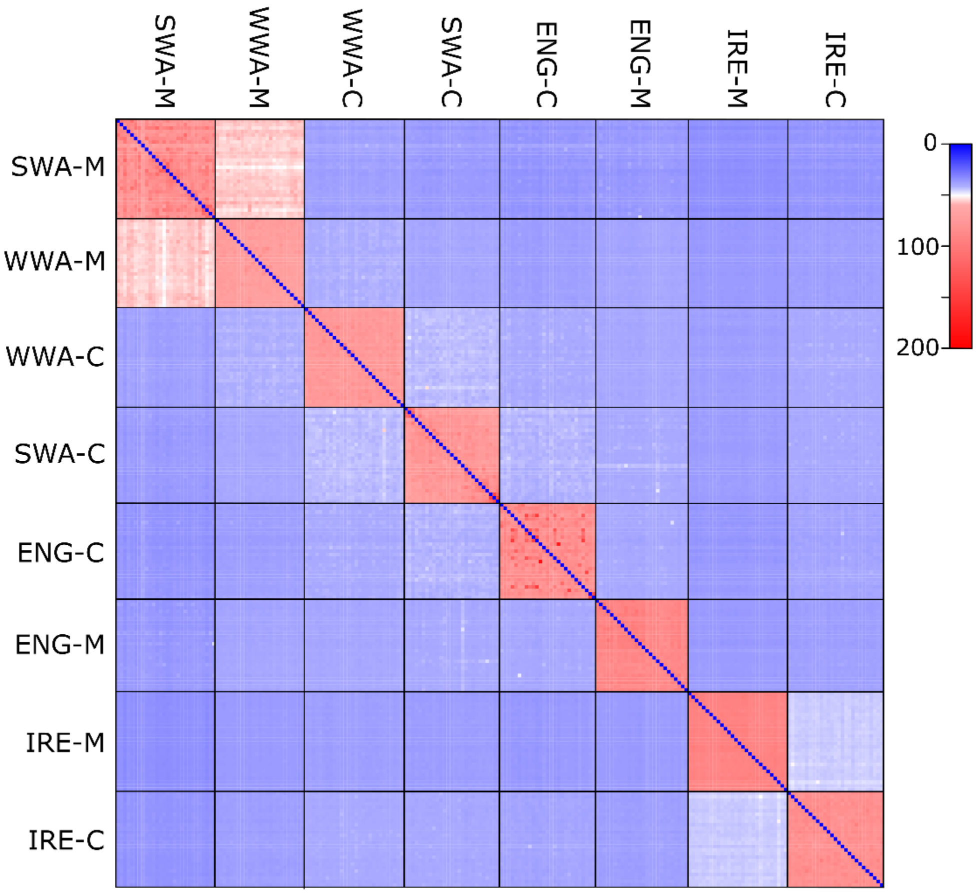

**Figure S2.** Treemix analysis with four migration edges. Colours as in Figure 1. The topology is almost identical to that produced by the SNPhylo analysis - the relationship of ENG-C and SWA-C to ENG-M was reversed.

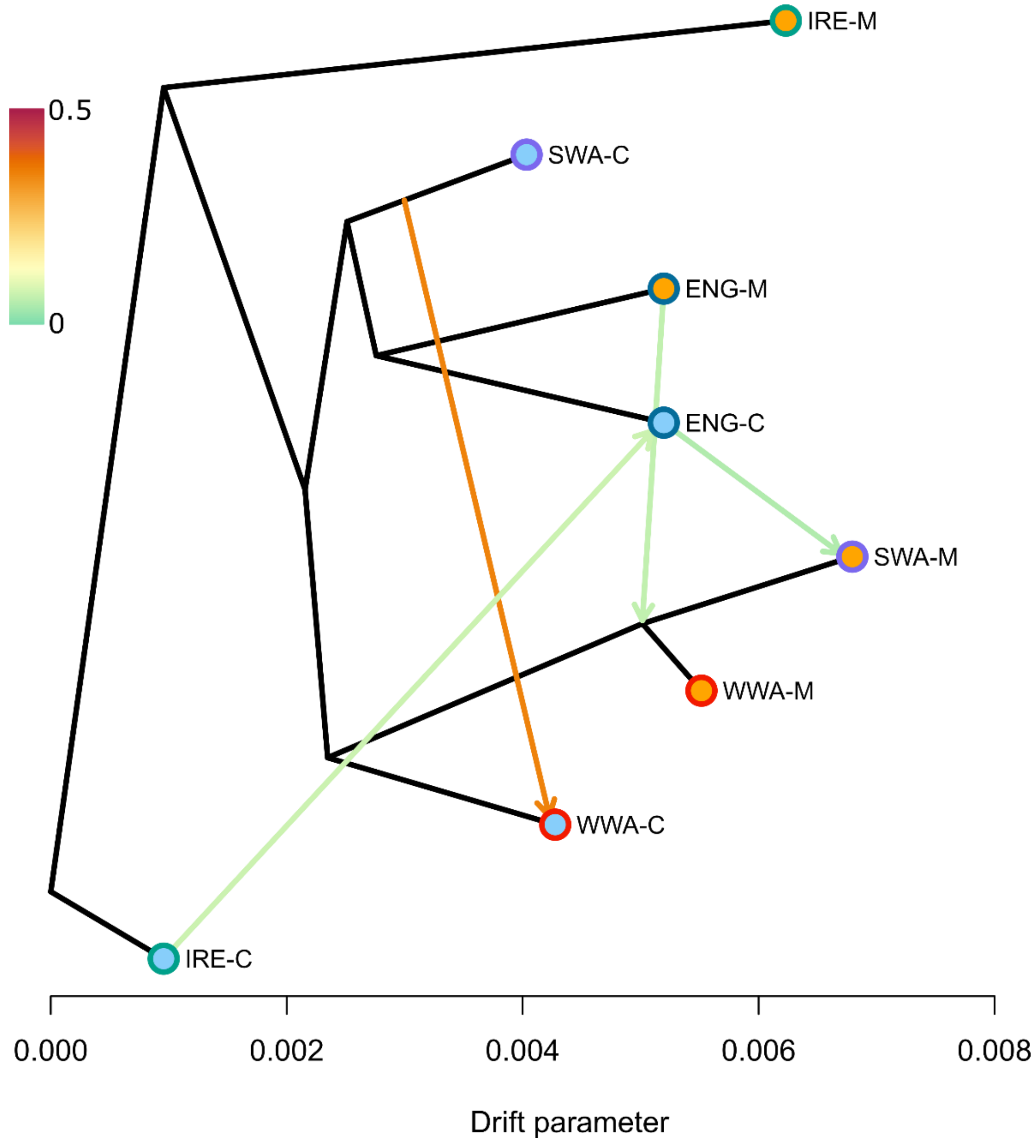

**Figure S3.** Tukey's test of association between number of outlier SNPs and scaffold length. 95% family-wise confidence levels

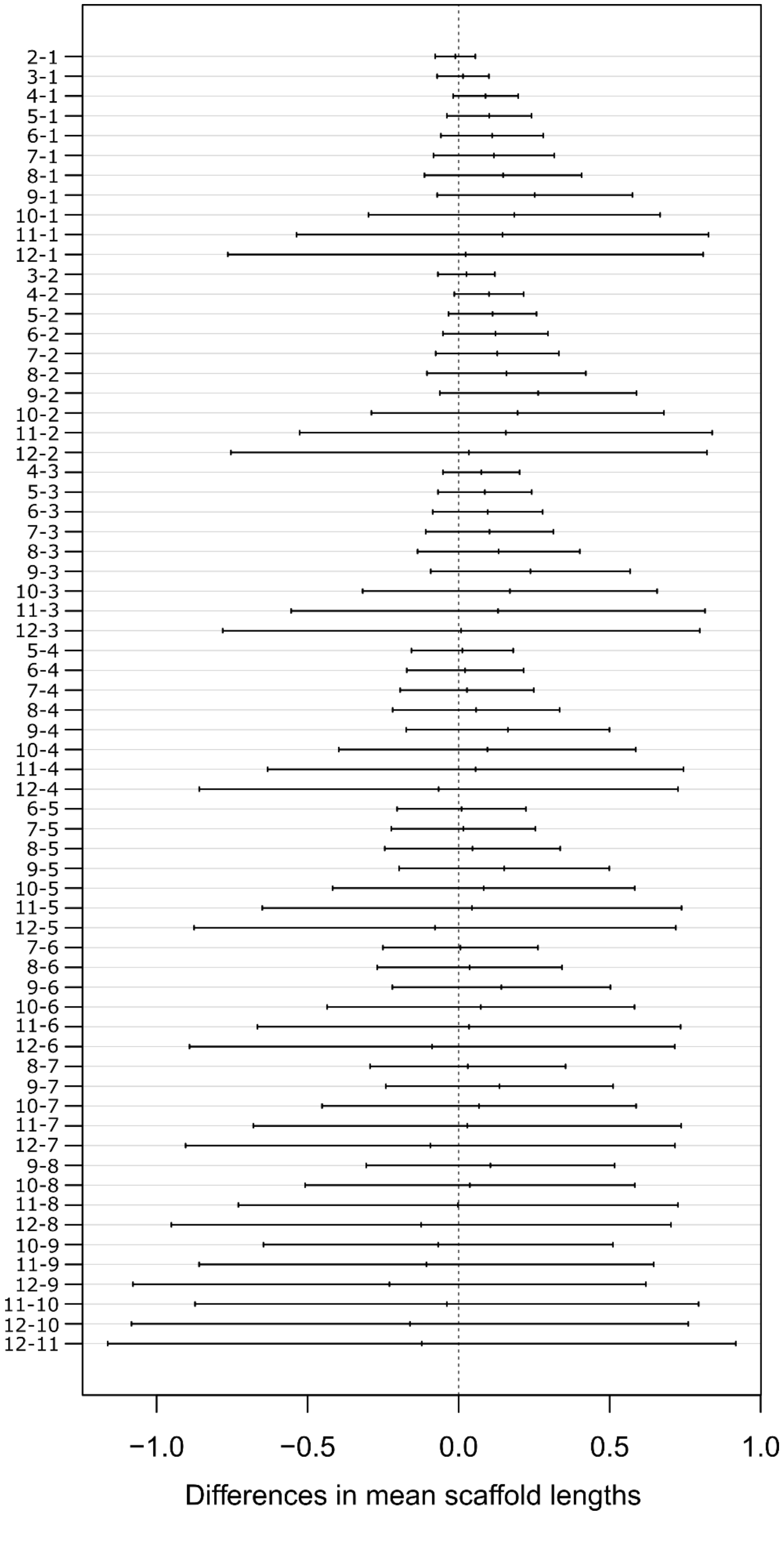

**Table S1.** Pairwise genetic differentiation ($F_{ST}$) between populations

| | **WWA-M** | **ENG-M** | **IRE-M** | **SWA-M** | **WWA-C** | **ENG-C** | **IRE-C** |
|---|---|---|---|---|---|---|---|
| **ENG-M** | 0.427 | - | - | - | - | - | - |
| **IRE-M** | 0.552 | 0.578 | - | - | - | - | - |
| **SWA-M** | 0.182 | 0.427 | 0.540 | - | - | - | - |
| **WWA-C** | 0.295 | 0.336 | 0.464 | 0.314 | - | - | - |
| **ENG-C** | 0.341 | 0.349 | 0.473 | 0.340 | 0.244 | - | - |
| **IRE-C** | 0.365 | 0.389 | 0.430 | 0.365 | 0.269 | 0.287 | - |
| **SWA-C** | 0.295 | 0.309 | 0.440 | 0.307 | 0.199 | 0.219 | 0.253 |

**Table S2**. Genetic diversity and Tajima's D for each population

| Population | Tajima's D | π | π *WRKY23* | π *RWP1* | π *PSD2* |
|---|---|---|---|---|---|
| WWA-M | 0.314 | 0.049 | 0.011 | 0.056 | 0.105 |
| WWA-C | 0.207 | 0.064 | 0.088 | 0.188 | 0.137 |
| SWA-M | 0.013 | 0.056 | 0.087 | 0.075 | 0.130 |
| SWA-C | 0.160 | 0.071 | 0.039 | 0.101 | 0.091 |
| ENG-M | 0.401 | 0.044 | 0.088 | 0.105 | 0.066 |
| ENG-C | 0.203 | 0.065 | 0.079 | 0.194 | 0.054 |
| IRE-M | 0.336 | 0.028 | 0.003 | 0.000 | 0.018 |
| IRE-C | 0.280 | 0.059 | 0.108 | 0.099 | 0.046 |

**Table S3.** Genome assembly statistics

| Assembly statistics | bp |
|---|---|
| n scaffolds | 269915 |
| n scaffolds > 5kb | 36751 |
| Largest contig | 151586 |
| Total length | 769407286 |
| GC (%) | 36.88 |
| N50 | 4660 |
| N75 | 2183 |
| L50 | 40594 |
| L75 | 101673 |
| N's per 100kbp | 982.85 |

| BUSCOs | n (%) |
|---|---|
| Complete BUSCOs | 1501 (64.5) |
| Complete and single-copy BUSCOs | 1368 (58.8) |
| Complete and duplicated BUSCOs | 133 (5.7) |
| Fragmented BUSCOs | 326 (14.0) |
| Missing BUSCOs | 499 (21.5) |

**Table S4** Gene ontology enrichment analysis for the 34 outlier scaffolds which were common to all ecotype pairs.

| GO ID | Term | Annotated | Significant | Expected | p-value |
|---|---|---|---|---|---|
| GO:0042938 | Dipeptide transport | 64 | 3 | 0.08 | 0.000058 |
| GO:0010337 | Regulation of salicylic acid metabolic process | 37 | 2 | 0.04 | 0.00087 |
| GO:0042939 | Tripeptide transport | 43 | 2 | 0.05 | 0.00117 |
| GO:0010310 | Regulation of hydrogen peroxide metabolic process | 53 | 2 | 0.06 | 0.00178 |
| GO:0000904 | Cell morphogenesis involved in differentiation | 634 | 4 | 0.75 | 0.00612 |
| GO:0051704 | Multi-organism process | 4313 | 11 | 5.09 | 0.00663 |
| GO:0009631 | Cold acclimation | 114 | 2 | 0.13 | 0.00794 |
| GO:0033036 | Macromolecule localization | 775 | 4 | 0.91 | 0.01224 |
| GO:0009860 | Pollen tube growth | 413 | 3 | 0.49 | 0.01233 |
| GO:0009617 | Response to bacterium | 1793 | 6 | 2.11 | 0.01603 |
| GO:0009932 | Cell tip growth | 472 | 3 | 0.56 | 0.01759 |
| GO:0048868 | Pollen tube development | 548 | 3 | 0.65 | 0.02598 |
| GO:0008104 | Protein localization | 552 | 3 | 0.65 | 0.02647 |
| GO:0048588 | Developmental cell growth | 569 | 3 | 0.67 | 0.02862 |
| GO:0051707 | Response to other organism | 3274 | 8 | 3.86 | 0.03032 |
| GO:0043207 | Response to external biotic stimulus | 3274 | 8 | 3.86 | 0.03032 |
| GO:0070727 | Cellular macromolecule localization | 582 | 3 | 0.69 | 0.03032 |
| GO:0009607 | Response to biotic stimulus | 3275 | 8 | 3.86 | 0.03037 |
| GO:0044706 | Multi-multicellular organism process | 710 | 3 | 0.84 | 0.04992 |
| GO:0009856 | Pollination | 710 | 3 | 0.84 | 0.04992 |

**Table S5.** Functional information for genes on 34 common outlier scaffolds

| Scaffold | Gene ID | Locus Identifier | Gene Symbol | Relevant function | Evidence | References |
|---|---|---|---|---|---|---|
| scaffold14499 | g46038 | AT2G02450 | LONG VEGETATIVE PHASE 1 (LOV1) | Cold | Regulates the plant cold response by positive regulation of the cold response genes COR15A and KIN1. Interacts with CBF/DREB1 genes (CBF2) | https://doi.org/10.1371/journal.pone.0000642 |
| scaffold00101 | g707 | AT4G25470 | C-REPEAT/DRE BINDING FACTOR 2 (CBF2) | Cold; Drought; Salt | Key role in cold tolerance and plays role in drought and salt tolerance. Also induced under cadmium stress in rice | https://doi.org/10.1093/jxb/err279 https://doi.org/10.1007/s11104-009-0116-9 https://doi.org/10.1093/jxb/erp300 |
| scaffold00735 | g4576 | AT5G17310 | UDP-GLUCOSE PYROPHOSPHORYLASE 2 (UGP2) | Cold; Roots | Affects root length expression changes under cold stress and heat shock in *A. thaliana* | https://doi.org/10.1093/pcp/pcp052 |
| scaffold12242 | g41152 | AT1G32060 | PHOSPHORIBULOKINASE (PRK) | Cold; Salt | Component of Calvin cycle. Expression inhibited by cold and salt stress in *Mesembryanthemum crystallinum.* | https://doi.org/10.1007/BF00028789 |
| scaffold17088 | g51384 | AT5G55190 | RAN GTPASE 3 (RAN3) | Cold; Salt | Expression increases in response to cold, salt and ABA stress, and *A. thaliana* mutants are sensitive to salt and ABA treatment. CBF2 expression levels were lower in a atran1/atran3 double mutant 10.1371/journal.pone.0154787 | https://doi.org/10.1371/journal.pone.0154787 |
| scaffold04408 | g19664 | AT5G54270 | LIGHT-HARVESTING CHLOROPHYLL B-BINDING PROTEIN 3 (LHCB3) | Drought; Inhibited by Zinc | Involved in stomatal response to abscisic acid and therefore important for drought stress response Zinc inhibits Photosystem II core complex. | https://doi.org/10.3390/ijms19030787 https://doi.org/10.1104/pp.66.6.1174 https://doi.org/10.1007/BF00042010 |
| scaffold09338 | g34010 | AT3G54140 | ARABIDOPSIS THALIANA PEPTIDE TRANSPORTER 1 (AtPTR1) | Heavy metal | Possible role in glutathione transport and cadmium tolerance. | https://doi.org/10.1016/j.febslet.2007.04.047 |
| scaffold00902 | g5465 | AT3G53000 | PHLOEM PROTEIN 2-A15 (PP2-A15) | Heavy metal | Protein specifically expressed in *Arabis paniculata* (hyper accumulator) in response to zinc stress. | https://doi.org/10.1016/j.chemosphere.2010.10.030 |
| scaffold00735 | g4578 | AT5G57190 | PHOSPHATIDYLSERINE DECARBOXYLASE 2 (PSD2) | Heavy metal | Located in tonoplast and produces phosphatidylethanolamine. In *S. cervisiae* is required for cadmium tolerance. Pseudomonas fluorescens produces phosphatidylethanolamine to avoid effects millimolar amounts of metals including Zinc | https://doi.org/10.1091/mbc.e09-06-0519 https://doi.org/10.1111/j.1574-6968.1995.tb07753.x |
| scaffold06199 | g25472 | AT2G47260 | WRKY DNA-BINDING PROTEIN 23 (WRKY23) | Heavy metal; Roots | WRKY23 regulates root development by affecting auxin distribution through the control of flavonol biosynthesis in *A. thaliana*. It increases quercetin root concentrations in overexpression mutants, whereas WRKY23 reduction of expression roots lack quercetin-rhamnoside-glucoside. Quecetin chelates metal ions can induce root growth in plants inhibited by Zinc. | https://doi.org/10.1073/pnas.1121134109 |
| scaffold00074 | g528 | AT4G29010 | ABNORMAL INFLORESCENCE MERISTEM (AIM1) | Heavy Metal; Salt | Key component of jasmonic acid (JA) synthesis and benzoic acid (BA) syntheses. JA is involved in both salt stress responses and heavy metal responses in plants. BA is known to be important for salt stress. | https://doi.org/10.1093/jxb/erw202 https://doi.org/10.1631/jzus.B1700191 |
| scaffold01631 | g8909 | AT5G33340 | CONSTITUTIVE DISEASE RESISTANCE 1 (CDR1) | Heavy Metal; Salt | Triggers expression of PR1 and PR2 which are part of salicylic acid pathway. Overexpression of PR1 and PR3 in tobacco plants leads enhanced tolerance to salt and heavy metals | https://doi.org/10.1038/sj.emboj.7600086. https://doi.org/10.1016/j.micres.2018.04.008 https://doi.org/10.1016/j.micres.2018.04.008 |

**Table S5.** Continued…

| Scaffold | Gene ID | Locus Identifier | Gene Symbol | Relevant function | Evidence | References |
|---|---|---|---|---|---|---|
| scaffold02487 | g12589 | AT5G41040 | REDUCED LEVELS OF WALL-BOUND PHENOLICS 1 (RWP1) | Heavy Metal; Salt | Required for suberin synthesis, lack of suberin in RWP1 mutants in *A. thaliana* leads to increased salt sensitivity. Suberin also contributes to blocking metal uptake from roots to shoots. | https://doi.org/10.1073/pnas.0905555106 https://doi.org/10.1371/journal.pgen.1000492 |
| scaffold06199 | g25474 | AT2G17030 | SKP1/ASK-INTERACTING PROTEIN 23 (SKIP23) | Heavy Metal; Salt | Differentially expressed under salt and mercury treatment in *Medicago trunculata* | https://doi.org/10.1007/s10142-015-0438-z |
| scaffold00284 | g1919 | AT3G57240 | BETA-1,3-GLUCANASE 3 (BG3) | Pathogen | Response to fungal pathogens | https://doi.org/10.1093/molbev/msm024 |
| scaffold06199 | g25471 | AT2G03200 | ATYPICAL ASPARTIC PROTEASE IN ROOTS 1 (ASPR1) | Roots | Involved in regulation of root development in *A. thaliana*. Overexpression leads to shorter primary roots and fewer lateral roots. Possible role in redox signalling in roots | https://doi.org/10.1093/jxb/erz059 |
| scaffold00463 | g3016 | AT2G31280 | CONSERVED PEPTIDE UPSTREAM OPEN READING FRAME 7 (CPuORF7) | Roots | Loss of function and overexpression mutants in *A. thanliana* show that LHL3 plays a positive role in xylem differentiation downstream of auxin | https://doi.org/10.1093/pcp/pct013 |
| scaffold00556 | g3575 | AT5G50010 | Basic helix-loop-helix 145 (SACL2) | Salt | Possible role in salt tolerance - quadruple T-DNA insertion mutant of the SAC51 family showed increased salt sensitivity in *A. thaliana* | https://doi.org/10.1111/tpj.14448 |
| scaffold00101 | g715 | AT4G23160 | CYSTEINE-RICH RLK (RECEPTOR-LIKE PROTEIN KINASE) 8 (CRK8) | Salt | Salt stress response. Germination of Crk8 knockout in *A. thaliana* is inhibited by NaCl | https://doi.org/10.1371/journal.pgen.1005373 |
| scaffold00902 | g5467 | AT3G53030 | SER/ARG-RICH PROTEIN KINASE 4 (SRPK4) | Salt | Phosphorylates SCL30 and RSp31. RSp31 undergoes increased splicing under salt stress. SCL30 constitutively phosphorylated by SRPK4. | https://doi.org/10.1186/1471-2164-15-431 |
| scaffold04926 | g21384 | AT3G54420 | HOMOLOG OF CARROT EP3-3 CHITINASE (EP3) | Wound | Highly expressed in roots. Response to wounding, but no increased expression under abiotic stress. | https://doi.org/10.1271/bbb.80837 |
| scaffold43030 | g91266 | AT1G23360 | 2-phytyl-1,4-naphthoquinone methyltransferase (MENG) | - | - | - |
| scaffold03268 | g15586 | AT3G19910 | RING/U-box superfamily protein (CTL18) | - | - | - |
| scaffold14742 | g46598 | AT1G01900 | Subtilisin-like serine protease (SBTI1.1) | - | - | - |
| scaffold02487 | g12592 | AT3G01990 | ACT DOMAIN REPEAT 6 (ACR6) | - | - | - |
| scaffold09338 | g34011 | AT5G01180 | ARABIDOPSIS THALIANA PEPTIDE TRANSPORTER 1 (AtPTR5) | - | - | - |
| scaffold00284 | g1918 | AT3G57270 | BETA-1,3-GLUCANASE 1 (BG1) | - | - | - |
| scaffold28068 | g70516 | AT5G23530 | CARBOXYESTERASE 18 (CXE18) | - | - | - |
| scaffold43030 | g91265 | AT1G23380 | KNOTTED1-LIKE HOMEOBOX GENE 6 (KNAT6) | - | - | - |
| scaffold04408 | g19665 | AT5G54280 | MYOSIN 2 (ATM2) | - | - | - |
| scaffold02103 | g11039 | AT3G07680 | P24 SUBFAMILY BETA 2 (P24BETA2) | - | - | - |
| scaffold00735 | g4577 | AT1G03050 | PHOSPHATIDYLINOSITOL BINDING CLATHRIN ASSEMBLY PROTEIN 5A (PICALM5A) | - | - | - |
| scaffold18761 | g54571 | AT3G18680 | PLASTID 55 UMP KINASE (PUMPKIN) | - | - | - |
| scaffold00284 | g1916 | ATCG00190 | RNA POLYMERASE SUBUNIT BETA (RPOB) | - | - | - |
| scaffold00074 | g527 | AT2G27520 | - | - | - | - |
| scaffold00074 | g529 | AT4G29000 | - | - | - | - |
| scaffold00074 | g530 | AT4G28990 | - | - | - | - |
| scaffold00101 | g713 | AT5G23400 | - | - | - | - |
| scaffold00101 | g714 | AT1G21280 | - | - | - | - |
| scaffold00101 | g718 | ATMG00710 | - | - | - | - |
| scaffold00101 | g720 | AT4G29090 | - | - | - | - |
| scaffold00284 | g1917 | AT1G74630 | - | - | - | - |
| scaffold00284 | g1921 | AT5G07400 | - | - | - | - |
| scaffold00284 | g1923 | AT4G00980 | - | - | - | - |

**Table S5.** Continued…

| Scaffold | Gene ID | Locus Identifier | Gene Symbol | Relevant function | Evidence | References |
|---|---|---|---|---|---|---|
| scaffold00383 | g2509 | AT3G01410 | - | - | - | - |
| scaffold00383 | g2516 | ATMG00860 | - | - | - | - |
| scaffold00383 | g2520 | AT4G13320 | - | - | - | - |
| scaffold00463 | g3015 | AT5G45190 | - | - | - | - |
| scaffold00463 | g3017 | AT1G06640 | - | - | - | - |
| scaffold00556 | g3570 | AT1G20400 | - | - | - | - |
| scaffold00556 | g3576 | AT5G51560 | - | - | - | - |
| scaffold00735 | g4575 | AT5G14720 | - | - | - | - |
| scaffold00902 | g5466 | AT2G36630 | - | - | - | - |
| scaffold01259 | g7215 | AT3G19184 | - | - | - | - |
| scaffold01631 | g8908 | AT2G07760 | - | - | - | - |
| scaffold01631 | g8910 | AT1G64830 | - | - | - | - |
| scaffold01631 | g8914 | AT5G09430 | - | - | - | - |
| scaffold02487 | g12590 | AT5G14080 | - | - | - | - |
| scaffold03268 | g15587 | AT5G08310 | - | - | - | - |
| scaffold04926 | g21383 | AT2G05642 | - | - | - | - |
| scaffold06199 | g25466 | AT5G17410 | - | - | - | - |
| scaffold06911 | g27490 | AT1G33710 | - | - | - | - |
| scaffold06911 | g27491 | AT1G16220 | - | - | - | - |
| scaffold09359 | g34066 | AT5G16210 | - | - | - | - |
| scaffold12242 | g41153 | AT5G45590 | - | - | - | - |
| scaffold28068 | g70515 | AT5G05180 | - | - | - | - |
| scaffold119902 | g154220 | AT5G19890 | - | - | - | - |
| scaffold168321 | g177089 | AT5G56190 | - | - | - | - |

**Table S6.** Population GPS coordinates

| | Latitude | Longitude |
|---|---|---|
| WWA-M | 52.331608 | -3.887207 |
| WWA-C | 52.394825 | -4.093914 |
| SWA-M | 51.635815 | -3.933929 |
| SWA-C | 51.566044 | -4.003153 |
| ENG-M | 51.256935 | -2.650950 |
| ENG-C | 51.323284 | -3.016996 |
| IRE-M | 52.034954 | -9.539131 |
| IRE-C | 52.128418 | -9.898941 |
